# An evolutionarily acquired microRNA shapes development of mammalian cortical projections

**DOI:** 10.1101/2020.09.08.286955

**Authors:** Jessica L Diaz, Verl B Siththanandan, Victoria Lu, Nicole Gonzalez-Nava, Lincoln Pasquina, Jessica L MacDonald, Mollie B Woodworth, Abduladir Ozkan, Ramesh Nair, Zihuai He, Vibhu Sahni, Peter Sarnow, Theo D Palmer, Jeffrey D Macklis, Suzanne Tharin

**Author notes:** These authors contributed equally (Jessica L Diaz, Verl B Siththanandan, Victoria Lu). Co-senior authors (Jeffrey D Macklis, Suzanne Tharin).

## Abstract

The corticospinal tract is unique to mammals and the corpus callosum is unique to placental mammals (eutherians). The emergence of these structures is thought to underpin the evolutionary acquisition of complex motor and cognitive skills. Corticospinal motor neurons (CSMN) and callosal projection neurons (CPN) are the archetypal projection neurons of the corticospinal tract and corpus callosum, respectively. Although a number of conserved transcriptional regulators of CSMN and CPN development have been identified in vertebrates, none are unique to mammals and most are co-expressed across multiple projection neuron subtypes. Here, we discover seventeen CSMN-enriched microRNAs (miRNAs), fifteen of which map to a single genomic cluster that is exclusive to eutherians. One of these, miR-409-3p, promotes CSMN subtype identity in part via repression of LMO4, a key transcriptional regulator of CPN development. *In vivo*, miR-409-3p is sufficient to convert deep-layer CPN into CSMN. This is the first demonstration of an evolutionarily acquired miRNA in eutherians that refines cortical projection neuron subtype development. Our findings implicate miRNAs in the eutherians’ increase in neuronal subtype and projection diversity, the anatomic underpinnings of their complex behavior.

**Significance Statement:** The mammalian central nervous system contains unique projections from the cerebral cortex thought to underpin complex motor and cognitive skills, including the corticospinal tract and corpus callosum. The neurons giving rise to these projections - corticospinal and callosal projection neurons - develop from the same progenitors, but acquire strikingly different fates. The broad evolutionary conservation of known genes controlling cortical projection neuron fates raises the question of how the more narrowly conserved corticospinal and callosal projections evolved. We identify a microRNA cluster selectively expressed by corticospinal projection neurons and exclusive to placental mammals. One of these microRNAs promotes corticospinal fate via regulation of the callosal gene LMO4, suggesting a mechanism whereby microRNA regulation during development promotes evolution of neuronal diversity.

## Introduction

The size and complexity of the mammalian brain dramatically increased with the evolution of placental mammals (eutherians). Eutherian evolutionary innovations included the consolidation of the motor cortex, the completion of the corticospinal tract, and the appearance of the corpus callosum (1–4). This evolution expanded both the overall number and the number of distinct subtypes of cortical projection neurons – the large, excitatory pyramidal neurons with axons that project long distances to deep, contralateral, or sub-cerebral structures.

The archetypal cortical projection neurons of the eutherian corticospinal tract and corpus callosum are, respectively, corticospinal motor neurons (CSMN) and callosal projection neurons (CPN). CSMN project from the motor cortex via the corticospinal tract to the spinal cord. They are centrally involved in the most skilled voluntary movement. By contrast, CPN project from the cerebral cortex via the corpus callosum to the contralateral cortex. They are involved in associative and integrative function (5). Despite their dramatically different projections and functions, CSMN and CPN are closely related in their development; CSMN and deep layer CPN are born at the same time and share common progenitors (5–8).

CSMN, CPN, and other cortical projection neurons develop through waves of neurogenesis, radial migration, and differentiation from radial glial progenitors of the ventricular zone and intermediate progenitor cells of the subventricular zone (9–12). The six-layered mammalian neocortex is generated in an inside-out fashion, with the earliest-born neurons populating the deepest layers, and the last-born neurons populating the most superficial layers (13). In the mouse, CSMN and a subset of CPN are generated around embryonic day 13.5 (e13.5), and both reside in the deep cortical layer V (Figure 1A). A larger subset of CPN is generated around e15.5, and it populates the superficial cortical layer(s) II/III (Figure 1A). A body of research from the last ~15 years has uncovered a set of key molecules, largely transcriptional regulators, that control cortical projection neuron development. These discoveries have been central to defining the current paradigm of combinatorial transcription factor controls (6–8).

**Figure 1.**
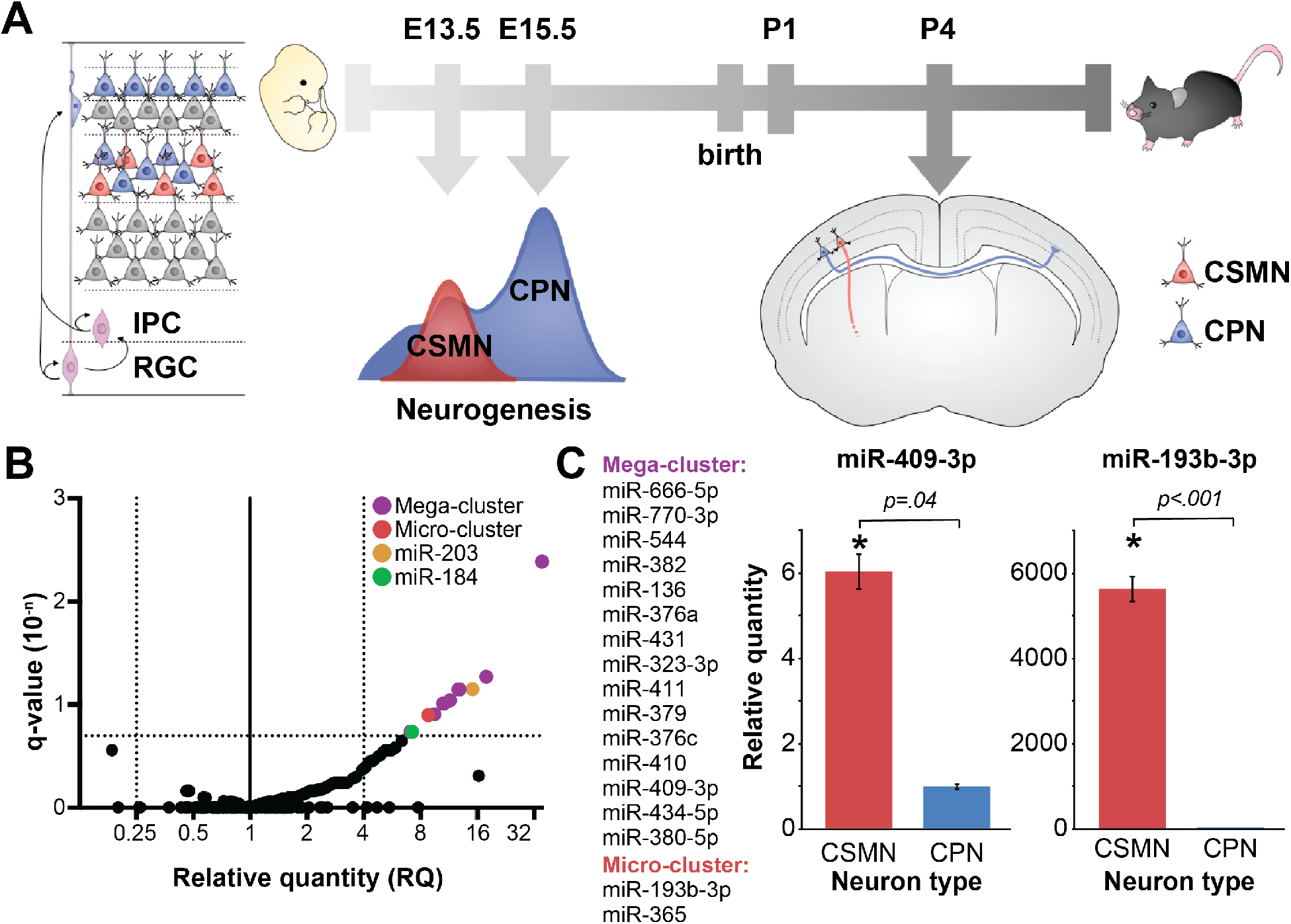
miRNAs are differentially expressed by CSMN vs. CPN during their development. (A) Schematic of CSMN (red) and CPN (blue) development in mice. (B) Volcano plot of differential miRNA expression by CSMN vs. CPN on P1, with fold change expressed as Relative Quantity (RQ) plotted against statistical significance expressed as False Discovery Rate-adjusted p-value (q-value), reveals 19 miRNA candidates enriched at least four-fold in CSMN relative to CPN with a q-value of less than 0.2 (colored dots). (C) Two confirmed miRNA clusters, comprising 17 differentially expressed candidate miRNAs, and representative results of independent validation via qPCR on P2. miR-409-3p from mega-cluster is 6-fold enriched (RQ=6) in CSMN vs. CPN on P2; miR193b-3p is 5630-fold enriched in CSMN vs. CPN on P2. Error bars represent SEM.

A number of transcription factors required for cortical projection neuron development have been identified as differentially expressed by CSMN, CPN, and other cortical projection neuron subtypes, and their roles delineated by functional analyses. For example, Fezf2 is a transcription factor required for specification of CSMN and other deep-layer subcerebral projection neuron subtypes (14, 15). Fezf2 acts as a selector gene, regulating the expression of neurotransmitter and axon guidance receptor genes in CSMN (15–20). SATB2, on the other hand, is a transcription factor required for CPN identity. SATB2 regulates the expression of a distinct set of downstream genes (21–23). Curiously, SATB2 is also required for early CSMN development, including extension of CSMN axons to the corticospinal tract (23). LIM domain Only 4 (LMO4) is a LIM-homeodomain transcription factor that is initially expressed by both CSMN and CPN, but becomes progressively restricted to CPN by late development (24). LMO4 is important for the establishment of CPN areal identity (25), and is also required for projection target diversity of CPN within motor cortex (26). LMO4, along with Fezf2, SATB2, and other developmental transcription factors, is co-expressed across multiple projection neuron subtypes early in their development, and is broadly conserved across non-eutherian vertebrates (6). This raises the question of what molecular mechanisms could underlie the projection neuron specializations acquired in eutherians.

Here, we provide evidence supporting the evolutionary emergence of a miRNA cluster that might underlie CSMN and CPN specializations acquired in eutherians. miRNAs are small non-coding RNAs that cooperatively repress multiple specific target genes post-transcriptionally (27). As such, they have the potential to regulate molecular programs that control or refine cellular identity. Following on previous findings that miRNAs play a critical role in early cortical development (28, 29), our studies reported here were designed to examine whether miRNAs control and/or refine development of CSMN and CPN in mice. Here, we identify that miR-409-3p is differentially expressed by developing CSMN vs. CPN, that it promotes CSMN fate in part through repression of the CPN-expressed transcriptional regulator LMO4, and that it is encoded on a cluster of miRNAs enriched in CSMN development that co-evolved with the motor cortex and corpus callosum. To our knowledge, this is the first report that miRNA repression of a specific transcriptional regulatory pathway shapes cortical projection neuron development and subtype identity.

## Results

### miRNAs are differentially expressed by CSMN and CPN during development

Multiple genes critical to cortical projection neuron development are differentially expressed by CSMN vs. CPN across eutherians, extensively studied in mice (6, 15, 22, 30–35). Our studies considered the possibility that this differential mRNA expression is in part controlled by miRNAs, which are known to coordinately regulate gene expression (27, 36). In a first series of experiments to address this question, we investigated whether miRNA expression is regulated in a neuron subtype-specific way in mice. We began by examining miRNA expression by CSMN vs. CPN at a critical developmental time point: on postnatal day one (P1), when these neurons express multiple lineage-restricted genes, but when the majority of their axons have not yet reached their targets (37).

CSMN and CPN were retrogradely labeled via ultrasound-guided nano-injection of fluorescent latex microspheres into their distal axon projections. Pure populations were obtained by fluorescence activated cell sorting (FACS), as previously described (30, 34, 38). miRNA expression was examined using TaqMan Low Density Arrays (TLDA), and analyzed using the comparative cycles to threshold (Ct) method (39). These experiments were designed to compare the Relative Quantity (RQ) of miRNA in one cell type (e.g. CSMN) relative to the other cell type (e.g. CPN), calculated as 2^−ΔΔCt^. The 2^−ΔΔCt^ method is particularly well suited to experiments in which there are limiting quantities of input RNA, and in which it is of greater biological relevance to compare the relative difference in expression between two samples than to determine the absolute transcript copy number, as in absolute quantification (2^−ΔC’t^) methods (39). The data are expressed in a volcano plot, depicting biological significance (fold-change) vs. statistical significance (False Discovery Rate-adjusted q values). From the 518 miRNAs investigated using these arrays, we identified 19 candidate miRNAs that are at least 4-fold enriched in CSMN vs. CPN under an FDR of 0.2 at P1 (Figure 1B). A complete list of candidates is found in Figure 1B-C. We examined the genomic organization of the 19 candidate CSMN-enriched miRNAs, and discovered that 15 of them map to a single, large miRNA cluster on mouse chromosome 12, the 12qF1 miRNA cluster. Two others comprise a micro-cluster on mouse chromosome 16. The remaining two candidates, miR-203 and miR-184, are individually encoded (Figure 1B). We independently validated at least one miRNA from each gene/cluster by qPCR. We purified independent samples of CSMN and CPN in biological triplicate on P2, and determined the RQ of each miRNA in CSMN relative to CPN using sno202 as a control. We confirmed that representative members of the mega and micro clusters are differentially expressed by CSMN vs. CPN on P2 (Figure 1C); on the other hand, miR-203 and miR-184 are equally expressed by CSMN vs. CPN on P2 (not shown), most likely representing false discoveries.

### miR-409-3p is enriched in CSMN, and represses the CPN transcriptional regulator LMO4

miRNAs often control developmental pathways by cooperatively repressing specific target genes. We carried out bioinformatic analyses to identify predicted targets of the differentially expressed miRNAs using the search tools miRanda (40–43), Targetscan (44–48), DIANALAB (49–51), and miRDB (52, 53), each driven by a slightly different target prediction algorithm. These analyses strongly and consistently predicted that one of the CSMN-enriched miRNAs, miR-409-3p, targets the known CPN transcriptional regulator LMO4. LMO4 is a LIM-homeodomain transcription factor that is starkly excluded from CSMN (30) and is important for CPN areal identity and projection target diversity (25, 26). Because miR-409-3p is strongly, consistently, and initially uniquely predicted to target an established CPN transcriptional regulator, we selected miR-409-3p from the group of CSMN-enriched miRNAs as a first candidate for functional analysis. Compared to >350 other predicted miR-409-3p targets, LMO4 is both highly ranked across all four target prediction algorithms, and well established to function in CPN development and CPN-vs.-CSMN distinction in particular, making it a logical choice for initial target analysis. Supplemental Figure 1 shows the top ten over-represented biological processes from a Gene Ontology (GO) analysis of predicted miR-409-3p targets. miR-409-3p is 6.8-fold enriched (RQ=6.8) in CSMN vs. CPN at P1, and it was independently confirmed to be 6-fold enriched in CSMN at P2 by qPCR (Figure 2A). Intriguingly, LMO4 is initially expressed by both CSMN and CPN, but becomes progressively restricted to CPN by early postnatal life (24) (Figure 2B), as would be predicted if miR-409-3p were repressing its expression in CSMN.

**Figure 2.**
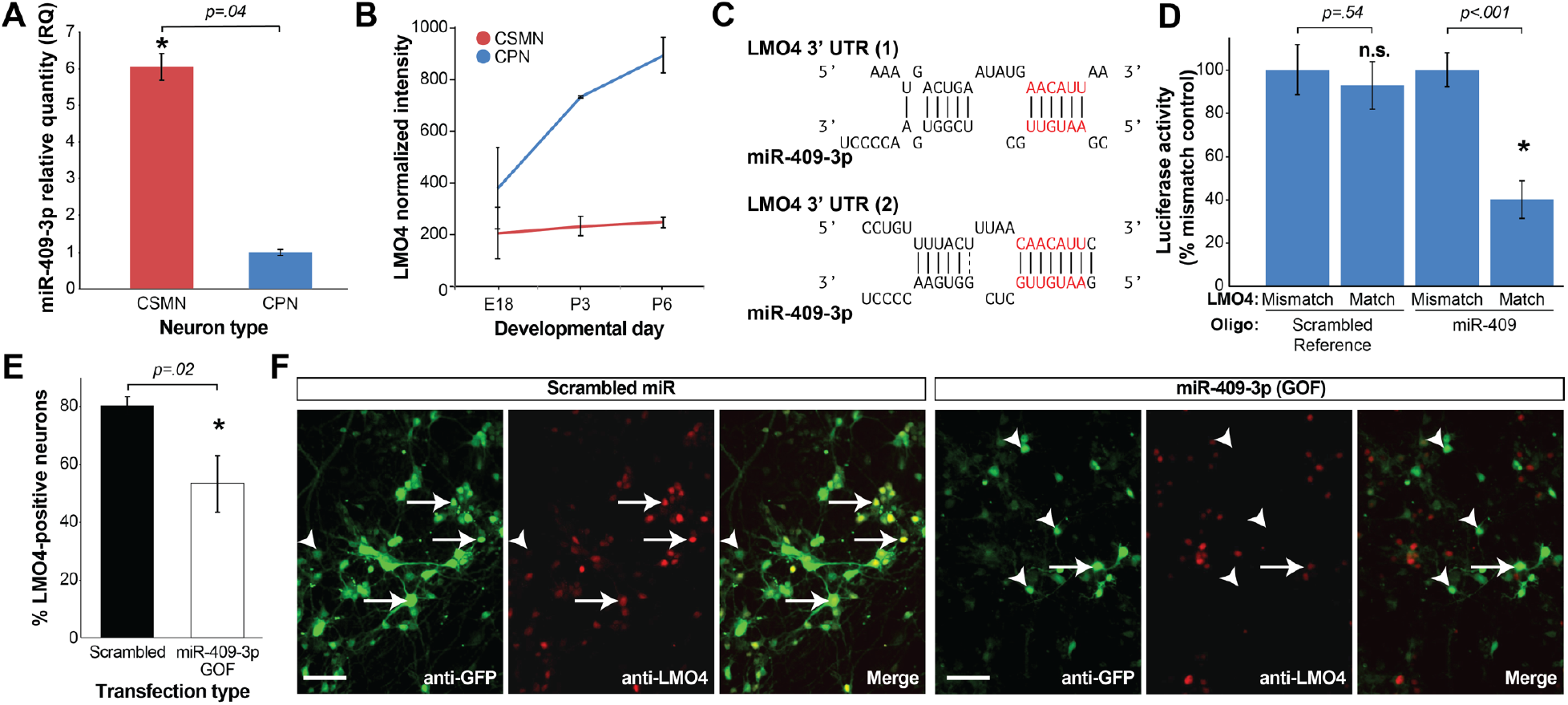
miR-409-3p is enriched in CSMN, and it represses the CPN-expressed and CSMN-excluded transcriptional regulator LMO4. (A) miR-409-3p is six-fold enriched in CSMN vs. CPN at P2 by qPCR. Error bars represent SEM. (B) LMO4 is enriched in CPN vs. CSMN in late embryonic and early postnatal life by microarray analysis. Error bars represent SEM. (C) Sequence alignments demonstrate that miR-409-3p is predicted to target two sites in the LMO4 3’ UTR. Seed sequence base-pairing is shown in red. (D) miR-409-3p oligonucleotides repress a LMO4 3’UTR luciferase reporter gene bearing wild-type, but not mismatch, miR-409-3p target sequences. Scrambled miRNA does not repress the LMO4 3’UTR luciferase reporter. Error bars represent SEM. * p<0.05 compared to mismatch control (E) Overexpression gain-of-function (GOF) of miR-409-3p in cultured embryonic cortical neurons results in decreased expression of LMO4, compared to scrambled control, by immunocytochemical analysis. Error bars represent SEM. * p<0.05 compared to scrambled control (F) Representative fluorescence micrographs illustrate reduction in LMO4 expression with overexpression of miR-409-3p. Scale bars, 50□m.

miR-409-3p is predicted to target two sites in the LMO4 3’ untranslated region (3’ UTR) (Figure 2C). To investigate whether miR-409-3p can use these sites to repress gene expression, we performed luciferase reporter gene assays in COS7 cells, as previously described (54, 55). We used LMO4 reporter vectors containing either wild-type or mutated (mismatch) miR-409-3p LMO4 target sites and their flanking 3’UTR sequences. We found that miR-409-3p oligonucleotides significantly and substantially repress LMO4 luciferase reporter gene expression by 60% with wild-type, but not mismatch, miR-409-3p target sequences (Figure 2D). Scrambled control miRNA oligonucleotides do not repress the LMO4 luciferase reporter gene.

To test whether this finding extends to endogenous LMO4 in mouse cortical neurons, we performed gain-of-function (GOF) overexpression experiments in cortical cultures, using lentiviral vectors expressing miR-409-3p and GFP. Similar vectors expressing a scrambled miRNA and GFP were used in parallel, and served as controls. Cultures of e14.5 cortical cells were transfected with these vectors, as described in Materials and Methods, and were examined for LMO4 expression by immunofluorescence labeling on day 7 in culture, a stage considered to be roughly equivalent to P1 *in vivo*. Double immunofluorescence with antibodies to LMO4 and GFP reveals that miR-409-3p overexpression leads to a reduction of the number of LMO4+ neurons in the targeted embryonic cortical cultures compared to the scrambled miRNA control (54% vs. 80% LMO4+ neurons; 33% decrease from control; Figure 2E,F), consistent with our luciferase reporter gene findings. miR-409-3p antisense loss-of-function (LOF) does not increase endogenous LMO4 expression in these cultures (Supplemental Figure 2), suggesting that additional miRNAs, possibly including six other CSMN-enriched 12qF1 cluster miRNAs (Figure 5B, C), redundantly repress LMO4 in cortical neurons, consistent with known mechanisms of cooperative miRNA repression (56). Collectively, the data indicate that miR-409-3p can repress expression of the CSMN-excluded and CPN-expressed transcriptional regulator LMO4 in cortical neurons, potentially thereby regulating subtype-specific cortical projection neuron development, distinction, and identity acquisition, CSMN development in particular.

### miR-409-3p promotes CSMN subtype identity, in part via LMO4 repression

To better understand the role of miR-409-3p in sculpting cortical projection neuron subtype identity, we carried out miR-409-3p overexpression GOF and antisense LOF experiments using cultured embryonic cortical neurons. Lentiviral vectors expressing miR-409-3p and GFP, and similar vectors expressing either an antisense miR-409-3p insert or a scrambled miRNA insert were used. Cultures of e14.5 cortical cells were transfected with these vectors as described in Materials and Methods, and were examined for marker expression by immunofluorescence labeling on day 7 in culture. To quantify the percent of CSMN in these cultures, we labeled with antibodies to CTIP2, a central CSMN/subcerebral projection neuron (SCPN) developmental control, and a canonical CSMN marker at high expression level at this developmental period, and to GFP, an indicator of transfected neurons. Relative to scrambled control, miR-409-3p transfected cultures (GOF) display a significant and substantial increase (11% increase in %CSMN; 68% change from control) in CTIP2+ neurons (Figure 3A, B). In contrast, miR-409-3p antisense (LOF) cultures display a decrease in CTIP2+ neurons (6.5% decrease in %CSMN; 40% change from control), although this is not statistically significant under a regression analysis taking all of the experiments (control/LOF/GOF/GOF+LMO4) into account (Figure 3A,B; Supplemental Table 1). ANOVA confirms that the means from these experiments differ from each other, consistent with our regression analysis (p = 0.0002). Our statistical analyses are summarized in Supplemental Table 1. Taken together, our results indicate that miR-409-3p favors CSMN development.

**Figure3.**
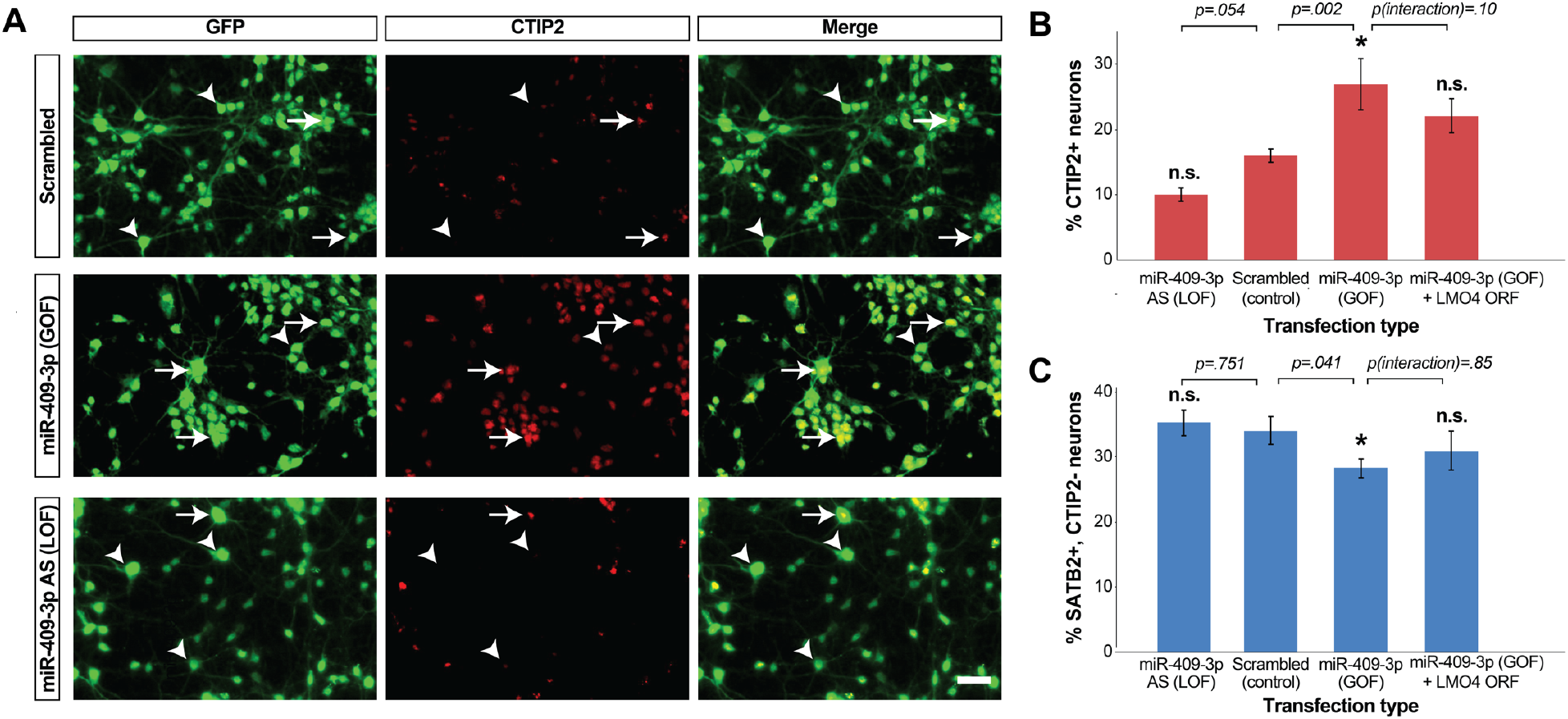
miR-409-3p promotes CSMN subtype identity, and inhibits CPN subtype identity, in part via LMO4 repression. (A) Representative fluorescence micrographs of embryonic cortical cultures illustrate an increase in the percent CTIP2+/GFP+ neurons (CSMN) with miR-409-3p GOF, and a decrease in the percent CTIP2+/GFP+ neurons with miR-409-3p LOF. Scale bar, 50μm. (B) miR-409-3p overexpression gain-of-function (GOF) increases the percent CTIP2+/GFP+ neurons (CSMN), and miR-409-3p antisense loss-of-function (LOF) decreases the percent CTIP2+/GFP+ neurons, compared to scrambled control in embryonic cortical cultures. Overexpression of the LMO4 open reading frame (ORF) reverses the miR-409-3p GOF phenotype in embryonic cortical cultures. (C) miR-409-3p GOF decreases the percent SATB2+/CTIP2−/GFP+ neurons (CPN) compared to scrambled controls in embryonic cortical cultures. Overexpression of the LMO4 ORF reverses the miR-409-3p GOF phenotype. Error bars represent SEM. *p<0.05 compared to scrambled control; n.s. not statistically significant compared to scrambled control; p(interaction) ~ modification of miR-409-3p GOF effect by LMO4 ORF.

To investigate whether miR-409-3p promotes CSMN development at least in part via repression of LMO4, we transfected cultures with lentiviral vectors expressing miR-409-3p, GFP, and the LMO4 open reading frame (ORF), directly assessing whether overexpression of the LMO4 ORF can suppress the miR-409-3p GOF phenotype. Unlike in miR-409-3p GOF, the percent CSMN in cultures overexpressing both miR-409-3p and LMO4 is statistically indistinguishable from that in scrambled control (p=0.074). This suggests suppression of the miR-409-3p GOF phenotype by overexpression of the LMO4 ORF. We then carried out a more stringent interaction analysis to evaluate how overexpression of the LMO4 ORF modifies the effect of miR-409-3p GOF. While we observed a relatively large magnitude of effect (−8.17%, standard error=4.7%), it is not statistically significant (p=0.1015) given the limited sample size. Taking all of the analyses into account, we interpret our findings as likely partial suppression, suggesting that miR-409-3p GOF acts in part through LMO4 repression. In separate experiments (Supplemental Figure 3), we have demonstrated that we do not observe an effect of LMO4 ORF overexpression on CTIP2 or SATB2 expression in these cultures, thus any interaction we observe does not represent an independent LMO4 overexpression phenotype. The lack of independent LMO4 overexpression phenotype (Supplemental Figure 3), together with the partial suppression of miR-409-3p GOF by LMO4, suggests that miR-409-3p affects the percent CSMN in these cultures via repression of multiple targets including LMO4, consistent with known multiple target mechanisms of miRNA action (Figure 5C) (56). Collectively, the data suggest that miR-409-3p favors CSMN development in part, but not exclusively, via repression of the CPN-expressed (and progressively CSMN-excluded) transcription factor LMO4.

### miR-409-3p inhibits CPN subtype identity

To quantify the percent CPN in miR-409-3p transfected cultures, we labeled with antibodies to SATB2, which is expressed by both deep and superficial layer CPN (21). Because SATB2 is co-expressed with CTIP2 by ~20-40% of CSMN during late embryonic and early postnatal development (23), we quantified the transfected CPN in our cultures by counting SATB2+/CTIP2−/GFP+ neurons, using triple label immunocytochemistry. Relative to scrambled control, the miR-409-3p transfected cultures (GOF) display a significant decrease in the percent CPN (6.2% decreased in %CPN; 18% change from control; Figure 3C) under a regression analysis taking all of the experiments (control/LOF/GOF/GOF+LMO4) into account. In contrast, the percent CPN in cultures transfected with the miR-409-3p antisense (LOF) lentivirus is statistically indistinguishable from that seen in the scrambled control transfections (0.9% increase in %CPN; 0.026% change from control; Figure 3C). To investigate whether miR-409-3p represses the adoption of CPN identity at least in part via repression of LMO4, we assessed whether overexpression of the LMO4 ORF can suppress the miR-409-3p GOF phenotype. Unlike in miR-409-3p GOF, the percent CPN in cultures overexpressing both miR-409-3p and LMO4 is statistically indistinguishable from that in scrambled control transfected cultures (p=0.33) (Figure 3C), suggesting suppression of the miR-409-3p GOF phenotype by overexpression of the LMO4 ORF. However, more stringent interaction analysis to evaluate how LMO4 modifies the effect of miR-409-3p GOF reveals a small magnitude, not statistically significant, effect (+0.72%, standard error=3.9%, p=0.85). While ANOVA does not show a significant overall mean difference across the experiments (control/LOF/GOF/GOF+LMO4; p = 0.0864), GOF is significantly different from control under a regression analysis (p = 0.041). Our statistical analyses are summarized in Supplemental Table 1. Taking all of our findings with respect to LMO4 overexpression into account, we interpret that miR-409-3p affects cortical projection neuron fate by acting through multiple targets, including LMO4. Collectively, the data suggest that miR-409-3p favors CSMN development in part, but not exclusively, at the expense of CPN development.

### miR-409-3p promotes corticospinal projection identity *in vivo*

To better understand the role of miR-409-3p in sculpting cortical projection identity, we carried out miR-409-3p overexpression GOF experiments *in vivo* via *in utero* electroporation. Plasmid vectors over-expressing miR-409-3p and tandem dimer Tomato (tdT), and similar vectors expressing a scrambled miRNA insert and tdT were used. Constructs were injected into the embryonic lateral ventricle on e13.5, during the peak of CSMN production and layer V CPN production. *In utero* electroporation was carried out as described in Materials and Methods. On e18.5, embryos were removed, fixed, and examined by immunofluorescence labeling both for marker expression and for axon projections. To quantify the percent of CSMN in these experiments, we again labeled with antibodies to CTIP2 and tdT, an indicator of transfected neurons. We controlled for electroporation efficiency between the control and experimental groups by calculating the number of CTIP2+ cells as a percentage of tdT+ (transfected) cells. Relative to scrambled control, miR-409-3p transfected cortices (GOF) have twice as many CTIP2+ neurons (22% increase in %CSMN; 100% change from control Figure 4A-C), indicating that miR-409-3p favors CSMN development *in vivo*, confirming our findings in primary culture. Importantly, qualitative anterograde visualization of transfected neurons reveals that miR-409-3p overexpression results in many more axons projecting along a subcerebral trajectory toward the internal capsule, and many fewer axons projecting medially toward the corpus callosum, compared to controls (Figure 4D-F). These results indicate that miR-409-3p not only promotes CSMN gene expression, but also corticospinal projection identity *in vivo*.

**Figure 4.**
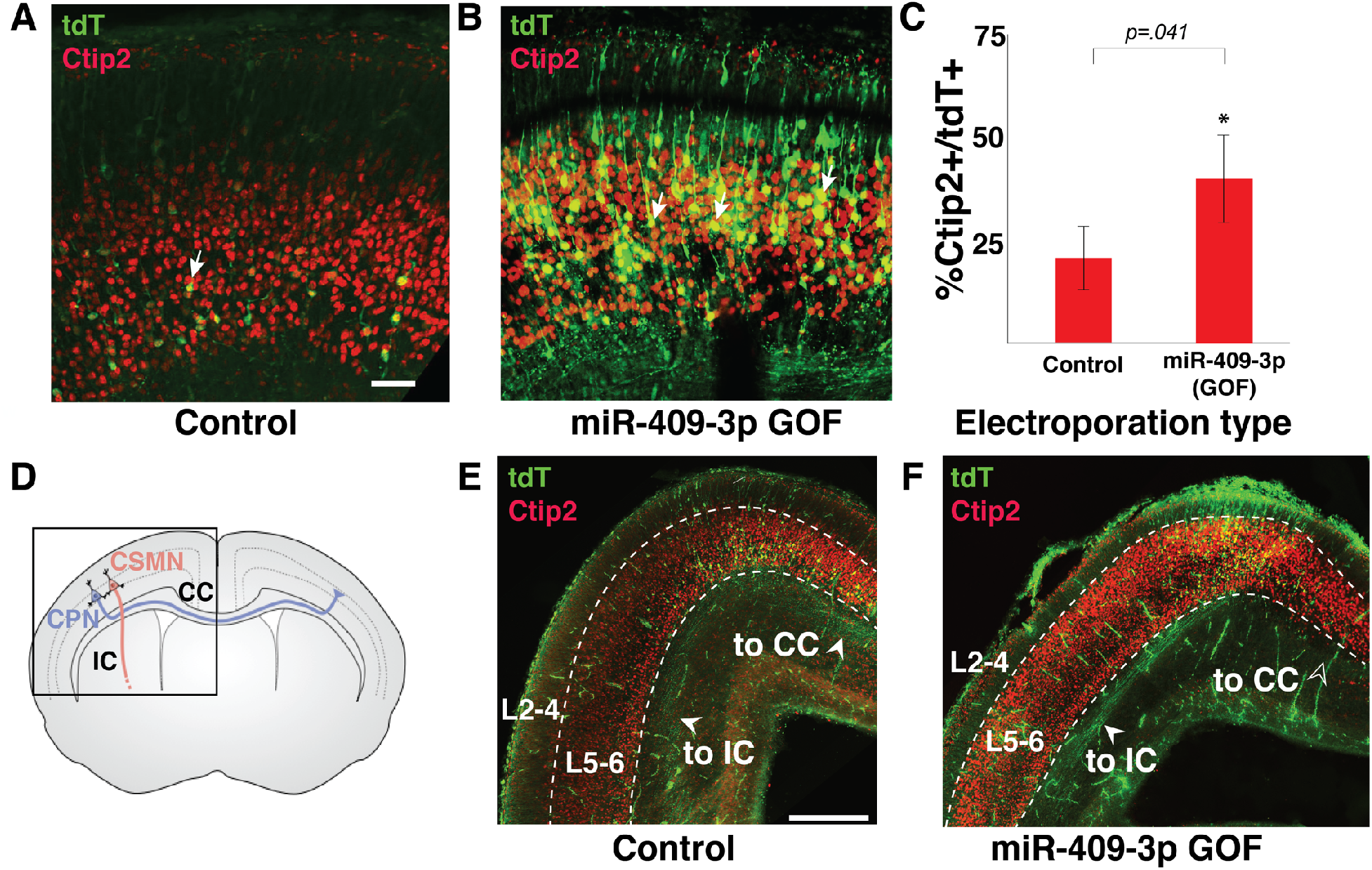
miR-409-3p promotes CSMN subtype identity and subcerebral axon trajectory *in vivo*. Representative fluorescence micrographs of e18.5 cortices electroporated at e13.5 illustrate an increase in the percent layer V CTIP2+/tdT+ neurons (CSMN, arrows) with miR-409-3p GOF (B), compared to scrambled control (A). Scale bar, 50μm. (C) miR-409-3p GOF results in a 100% increase (43.3% from 21.5%) in CTIP2+/tdT+ neurons (CSMN), compared to scrambled control *in vivo*. Error bars represent SEM. (D) Schematic of layer V CSMN (red) projecting sub-cerebrally via the internal capsule (IC) and CPN (blue) projecting inter-hemispherically via the corpus callosum (CC). (E-F) Representative coronal fluorescence micrographs of e18.5 brains electroporated at e13.5 illustrate many more axons projecting sub-cerebrally via the internal capsule (IC) and very few apparent axons projecting inter-hemispherically via the corpus callosum (CC) in miR-409-3 GOF (F) compared to in scrambled control (E). Scale bar, 500μm

### The 12qF1 miRNA cluster, miR-409-3p LMO4 target site 2, and the motor cortex and corpus callosum co-appeared with the evolution of eutherians

The clustered miRNAs appeared at the 12qF1 locus with the evolution of eutherians (Figure 5A), and are absent from this locus in marsupials, monotremes, and other vertebrates (57). While miR-409-3p LMO4 target site 1 is broadly conserved across vertebrates, site 2 is notably excluded from the genomes of marsupial mammals (e.g. koala) and is absent from all characterized chicken LMO4 mRNAs (Figure 5B). Clustered miRNAs have been shown to cooperatively repress the same gene, as well as interacting genes within a pathway (58). Updated bioinformatic algorithms, consistently predicting that miR-409-3p represses LMO4, now predict that six other CSMN-enriched 12qF1 cluster miRNAs also repress LMO4 (Figure 5B,C) (41, 42, 44–48, 51–53). Compellingly, all of these miRNAs are predicted to target sites within a portion of the LMO4 3’UTR that appears to be well conserved only among eutherians (Figure 5B). The motor cortex as a discrete areal region and the corpus callosum are, like the 12qF1 cluster miRNAs, evolutionary innovations of eutherians. The evolutionary relationship between the 12qF1 miRNAs, the motor cortex, and the corpus callosum further supports a role for these developmentally regulated miRNAs in the evolution of the archetypal projection neurons of these two structures: CSMN and CPN.

**Figure 5.**
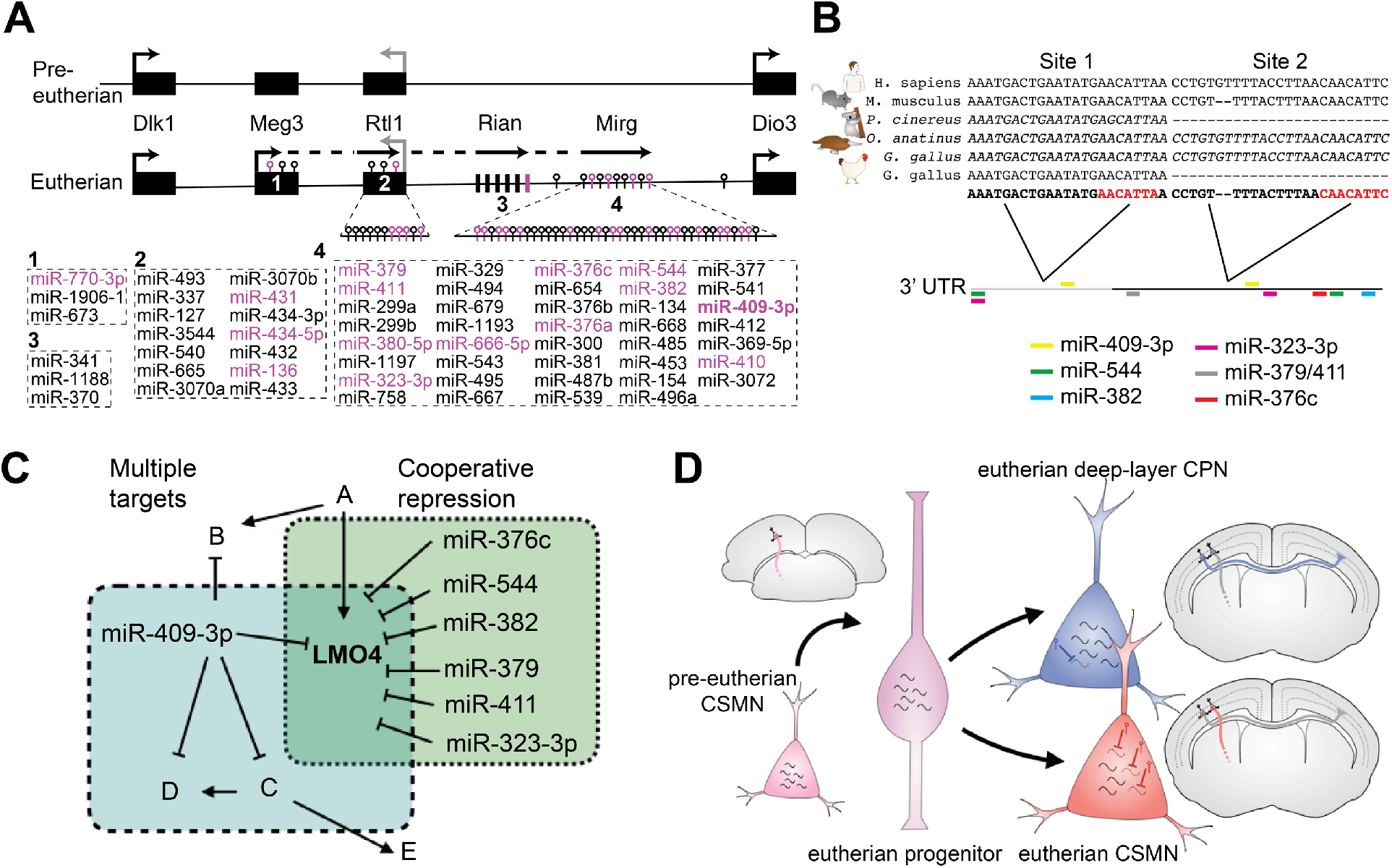
CSMN-enriched miRNAs are encoded at a genomic cluster that co-evolved with motor cortex and corpus callosum. (A) Schematic of the mouse 12qF1 locus highlighting miRNAs identified as enriched in CSMN (magenta). Meg3, anti-Rtl1, Rian, and Mirg are incompletely characterized genes encoding long RNAs that give rise to the eutherian-specific clustered 12qF1 microRNAs. Dlk1, Rtl1, and Dio3 are protein-coding genes at 12qF1 that are conserved in pre-eutherian mammals. (B) Schematic of the vertebrate LMO4 3’UTR depicts that the proximal portion (grey) is well-conserved among characterized vertebrate LMO4 mRNAs, whereas the distal portion of the eutherian LMO4 3’UTR (black) is absent from all characterized chicken LMO4 mRNAs and all marsupial LMO4 genes. The positions of predicted CSMN-enriched 12qF1 miRNA target sites, concentrated in the distal/eutherian portion of the LMO4 3’ UTR, are indicated by colored bars. Multiple sequence alignments illustrate that mir-409-3p site 1 is well conserved among vertebrates, whereas site 2 appears to be well conserved only among eutherians. Predicted mRNAs are italicized; characterized mRNAs are not italicized. (C) Individual miRNAs repress multiple targets, and clustered miRNAs cooperatively repress shared targets. Six CSMN-enriched 12qF1 cluster miRNAs, in addition to miR-409-3p, are predicted to cooperatively repress LMO4. (D) Model depicting deep layer CPN and eutherian CSMN derived from ancestral pre-eutherian CSMN via expansion of gene expression to generate CPN as a new projection neuron subtype, and pruning of this expansion in eutherian CSMN via miRNA-mediated repression of gene expression.

## Discussion

CSMN and CPN of layer V arise from a common progenitor pool, yet they acquire the highly divergent neuronal fates, projection identities, and circuit connectivities required for complex motor cortex output via the corticospinal tract, and expanded interhemispheric communication and integration via the corpus callosum, respectively. miRNAs have been implicated in early cortical development (28, 29), in production of cortical progenitors (59–77), in cortical neuronal migration (56, 66, 76, 78–83), and in the timing of neocortical layer production and progenitor competence (84). A mechanism of miRNAs controlling projection neuron fate specification in the nematode *C. elegans* has been demonstrated (85), however, no such mechanism yet has been demonstrated in mammals. A role for miRNAs in the development of cortical projection neuron subtypes has recently been postulated based on studies of miRNA-mRNA interaction networks during human cortical development (86). Curiously, that study did not identify any miRNAs to be enriched in newborn deep-layer neurons (which include multiple distinct, interspersed subtypes, including both CSMN and CPN, perhaps explaining the lack of identification), and identified a non-overlapping set to ours in maturing deep layer neurons. Species, developmental stage, and experimental approach differences most likely account for this. Here, we identify and functionally validate that miR-409-3p 1) is differentially expressed by developing CSMN vs. CPN, 2) promotes CSMN development in part through repression of the transcriptional regulator of CPN development LMO4, and 3) is encoded on the 12qF1 cluster of miRNAs enriched in CSMN development, which co-evolved with the motor cortex and corpus callosum.

LMO4 is a LIM-homeodomain transcription factor that is initially expressed by both CSMN and CPN, but becomes progressively restricted to CPN by late development (24), when it is required for the acquisition of motor CPN identity and projection complexity (25, 26). LMO4 was recently shown to permit co-expression of SATB2 and CTIP2 in two subclasses of somatosensory projection neurons (87), underscoring the importance of LMO4 repression to the acquisition of CSMN identity. We have shown that miR-409-3p represses LMO4 to favor CSMN development. Six other CSMN-enriched 12qF1 cluster miRNAs are also predicted to repress LMO4 (Figure 5B,C) (41, 42, 44–48, 51–53). Cooperative repression of a shared target by clustered miRNAs, well documented in miRNA biology (56, 58), is consistent with our finding that miR-409-3p LOF is not alone sufficient to de-repress LMO4 in cortical neurons even though elevated miR-409-3p is sufficient to repress LMO4. The potential role(s) of these additional miRNAs in shaping cortical projection neuron identity are not known. Collectively, these data suggest a model whereby the miRNAs of the 12qF1 cluster cooperatively repress a key CPN transcriptional regulator to promote and refine CSMN development and acquisition of identity.

Interestingly, LMO4 has been shown to bind directly to the cytoplasmic tail of Neogenin, an axon guidance receptor of the Deleted in Colorectal Cancer (DCC) family. LMO4-Neogenin binding is required to transduce repulsive signaling from RGM A through Neogenin in embryonic cortical neurons (88), raising the possibility that miR-409-3p and possibly other 12qF1 cluster miRNAs regulate CSMN axon guidance at least in part via repression of LMO4 during the development of CSMN projection identity. In support of our findings, an independent Gene Ontology (GO) analysis of predicted targets all of the 12qF1 cluster miRNAs previously revealed an over-representation of axon guidance genes (57). Future analyses could determine whether a subset of the 12qF1 cluster miRNAs repress specific axon guidance genes in CSMN to control projection identity, as the GO analysis suggests.

The partial rescue by LMO4 of the miR-409-3p GOF phenotype, the failure of LMO4 over-expression to phenocopy the miR-409-3p LOF phenotypes, and the decrease in proportion of CSMN despite no change in LMO4 in miR-409 LOF all suggest that miR-409-3p has additional target(s) relevant to CSMN and CPN development beyond LMO4. This is consistent with known multiple target mechanisms of miRNA action (Figure 5C) (27, 36), including during earlier aspects of neocortical development (56). Notably absent from even exhaustive lists of predicted miR-409-3p targets is the canonical CPN control gene SATB2. Our observations that miR-409-3p LOF alone may impair the adoption of CSMN fate, but does not result in a corresponding increased adoption of CPN fate or increase in LMO4 positive neurons, suggests that other 12qF1 microRNAs continue to impair the adoption of CPN fate, via repression of targets including, but not limited to, LMO4. The latter is consistent with known cooperative repression mechanisms of miRNA action (Figure 5C) (56, 58). Taken together, these findings support the provocative hypothesis that miR-409-3p, possibly in cooperation with other 12qF1 cluster miRNAs, refines CSMN fate by regulating novel cortical projection neuron genes that have yet to be identified and/or characterized.

This is the first description of a specific role for miR-409-3p in CSMN development, which we have functionally validated using primary cultures and *in vivo*. Additional *in vivo* functional studies, including germline and conditional deletion studies, of the roles of miR-409-3p and other 12qF1 miRNAs and long noncoding RNAs (lncRNAs) will be important avenues for future study. The 12qF1 lncRNA Meg3 was recently shown to play a critical role in lower motor neuron (LMN) development (89), and it is also enriched in developing CSMN (90). Taken together with our findings here, this suggests a greater role for the 12qF1 noncoding RNAs in the development of the corticospinal circuitry.

Studying the development of these archetypal projection neurons in mice could have important clinical implications in humans. CSMN, along with LMN, degenerate in amyotrophic lateral sclerosis (ALS) (91–93). Interestingly, miR-409-3p and a second 12qF1 cluster miRNA (miR-495-3p) have recently been implicated in the degeneration of embryonic stem cell-derived LMN bearing a mutation specific to a juvenile-onset form of ALS (94). Given the role of the 12qF1 lncRNAs in LMN development discovered recently (89), and the key role of miR-409-3p in CSMN development discovered here, it would be interesting to investigate whether miR-409-3p and other 12qF1 noncoding RNAs also play roles in the pathogenesis of ALS in CSMN.

We posit that eutherian CSMN and deep layer CPN are derived from ancestral pre-eutherian CSMN via at least two steps: an expansion of gene expression required to generate deep layer CPN as a new projection neuron subtype, combined with a pruning or refinement of this expansion in CSMN via miRNA-mediated repression (Figure 5D). The 12qF1 cluster is exclusive to eutherian mammals. In support of our model, we find that miR-409-3p LMO4 target site 2 is absent from marsupial LMO4 genes, and from all characterized chicken LMO4 mRNAs (Figure 5B). The 12qF1 cluster co-appeared during evolution with the motor cortex and the corpus callosum. Transcriptional control of the 12qF1 cluster, at least in skeletal and cardiac muscle, is via the MEF2A transcription factor. MEF2A is expressed in embryonic mouse cortex beginning at ~e13.5, the peak of CSMN and layer V CPN production (6, 7, 95). Genome-wide epigenetic analysis of the MEF2A cistrome in primary cortical cultures reveals widespread localization to enhancer elements of neurons, suggesting a possible role in neuronal lineage specification (96). Our model predicts that loss of 12qF1 cluster miRNA expression would result in a phenotype with dysgenesis of CSMN, potentially of their corticofugal pathfinding in particular (rather than an acallosal phenotype). Consistent with this prediction, MEF2A/D double knockout mice have significant deficits with rotarod motor control testing, but no other behavioral deficits reported (97), suggesting a relatively specific abnormality of corticospinal motor function.

miRNAs have been proposed to provide the cerebral cortex with “evolvability” (98, 99). An elegant example is the recent discovery that the great ape-specific miRNA miR-2115 represses the conserved cell cycle regulator ORC4 to control RCG proliferation during cortical development (86). The canonical transcription factors required for cortical projection neuron fate specification, including FEZF2, CITP2, SATB2, and LMO4, are broadly conserved across vertebrates (6, 8). Recent work also suggests that an intrinsic map of interhemispheric connections is conserved across mammals (100). Yet the motor cortex, fully elaborated corticospinal tract, and corpus callosum are exclusive to eutherian mammals. The 12qF1 miRNA cluster appeared with the emergence of eutherians, and axon guidance genes are enriched among its predicted targets. We have identified enrichment of a subset of 12qF1 miRNAs during CSMN development. We have further identified that at least one 12qF1 cluster miRNA, miR-409-3p, does indeed regulate LMO4, a transcription factor required for cortical projection neuron subtype specification and establishment of correct identity and projection diversity, and can promote CSMN fate. Taken together, our results indicate a central function of projection neuron subtype-specific, developmentally regulated miRNAs in sculpting and refining the specific identities of the expanded repertoire of cortical projection neurons of eutherian mammals.

## Acknowledgements

We thank Laure Aurelian for scientific discussions and for critically reading the manuscript. We thank members of the JDM, TDP, PS, and ST labs for scientific discussions and helpful suggestions. This work was supported by grants from the NIH (K08 NS091531), AOSpine North America (Young Investigator Research Grant Award), a Stanford McCormick Faculty Award, and a Stanford Maternal and Child Health Research Institute Pilot Award to ST. ST is a Tashia and John Morgridge Endowed Faculty Scholar in Pediatric Translational Medicine of the Stanford Maternal and Child Health Research Institute. This work was also supported by grants from the NIH (R01s NS045523 and NS075672, with additional infrastructure supported by NS041590, NS049553, and DP1 NS106665), the ALS Association, and the Travis Roy Foundation to JDM, by a grant from the NIH (R01 AI069000) to PS, and by grants from the NIH (1R01MH108660-01, 1R01MH108659-01, and R21NS096447) to TDP. JLD was supported by the comparative medicine biosciences training program (5T32OD011121-12). We acknowledge the Stanford Neuroscience Microscopy Service, supported by NIH NS069375. We gratefully acknowledge Michelle Monje for access to the electroporator and Paul Buckmaster for use of the sliding microtome.

## Author Contributions

ST and JDM conceived the project; ST, JDM, JLD, TDP, PS, and VBS designed the experiments; VBS, JLD, ST, LP, JLM, AO, and VS performed the experiments; VBS, VL, JLD, ZH, ST, and JDM analyzed and interpreted the data; NG, VL, and ST performed the bioinformatic analyses; ST, JLD, VBS, JLD, and MBW made the figures; ST, TDP, PS, and JDM synthesized and integrated the findings and wrote and revised the paper.

## Materials and Methods

### CSMN and CPN purification

CSMN and CPN were purified from C57BL/6J mice as previously described (15, 30, 34, 38). Briefly, CSMN were retrogradely labeled at P0 from the cerebral peduncle or at P1 from either the cerebral peduncle or spinal cord between C1 and C2 vertebrae under ultrasound microscopic guidance. At P1 or P2, neuron bodies in the motor cortex were isolated by microdissection of deep cortical layers, and dissociated to a single cell suspension by papain digestion and mechanical trituration. CPN were retrogradely labeled on P0 or P1 by injection of green fluorescent latex microspheres into the contralateral hemisphere under ultrasound microscopic guidance. On P1 or P2, labeled cortex was microdissected and dissociated to a single cell suspension by papain digestion and mechanical trituration. Neurons in suspension were FACS-purified into RNAlater (Life Technologies) using a FACSVantage sorter (BD), and purified labeled neuron cell bodies were then frozen at −80°C until RNA purification.

### RNA purification

microRNA was extracted from purified projection neuron cell bodies using the Ambion miRVana microRNA isolation kit (Ambion, Austin, TX) according to the manufacturer’s instructions. RNA quality was analyzed on a BioAnalyzer 2100 (Agilent), and confirmed to be excellent.

### Differential miRNA expression analysis

Comprehensive differential expression of 518 mouse miRNAs by CSMN and CPN at P1 was quantified using TaqMan Low Density Arrays (TLDA rodent miRNA v2.0, Applied Biosystems, Carlsbad, CA) at the Dana Farber Cancer Institute Molecular Diagnostics Laboratory (Boston, MA). The TLDA rodent miRNA v2.0 A and B cards correspond to Sanger miRbase version 10. The A card comprises well characterized miRNAs, whereas the B card comprises predicted miRNAs. qPCR data were analyzed using the comparative CT method (39), whereby miRNA expression is quantified based upon the number of PCR cycles required to reach detection threshold, normalized against the geometric mean of three endogenous control mouse miRNAs (sno135, sno202, U6). Relative Quantity (RQ) was calculated for each of the miRNAs, with RQ = 2^−□□Ct^, providing a measure of the fold difference in miRNA expression by one cell type vs. the other. We performed experiments in biological triplicate. To correct for multiple testing, False Discovery rate (FDR)-adjusted p-values (q-values) were calculated according to the method of Benjamini and Hochberg (101). Candidate miRNAs were independently validated via qPCR from purified P2 CSMN and CPN using sno202 as a control. We performed these experiments in biological triplicate.

### miRNA target prediction

We searched for predicted miRNA targets using the search tools miRanda (40–43), Targetscan (44–48), DIANALAB (49–51), and miRDB (52, 53).

### Luciferase assays

Luciferase reporter assays were performed using the Dual-Glo Luciferase Assay System (Promega), pmir-GLO based reporter constructs, and microRNA oligonucleotides (Horizon Discovery) according to the manufacturer’s instructions, as previously described (54, 55). Briefly, COS7 cells (10^4^/well) were seeded in a white 96-well plate. The following day, pmir-GLO reporter-miRNA oligo-DharmaFECT Duo (Dharmacon) transfection mixtures were prepared. The media from the 96-well plate was replaced with the transfection mixture, and the plate was incubated overnight. Firefly and renilla luciferase reporter fluorescence was read using a Tecan Infinite M1000 (Stanford High-Throughput Bioscience Center Core Facility). The ratio of firefly to renilla fluorescence was calculated for each well. Averages were compared for triplicates of each condition. Match reporter vectors contained the two wild-type predicted miR-409-3p seed regions (CAACATT) with 30bp of flanking LMO4 3’ UTR on either side of each. Mismatch reporter vectors were identical to match reporters except that the seed sequences were replaced by GGGGGGG. Additional negative controls using empty reporter vectors and scrambled control oligos, and positive controls using pmir-GLO miR21 reporter match/mismatch vectors and miR21 oligos, were performed. The experiments were replicated in n=5 independent cultures.

### Lentivirus vectors

Lentivirus vectors were modified from the pSicoR backbone (102), a gift from Tyler Jacks (Addgene plasmid # 11579). Expression of miRNA was under direction of the strong U6 promoter. miRNA inserts were either: miR-409-3p (gain of function: GAATGTTGCTCGGTGAACCCCTTTTTT), antisense miR-409-3p (loss of function: AGGGGTTCACCGAGCAACATTCTTTTT), or scrambled (control, CCTAAGGTTAAGTCGCCCTCGCTCCGAGGGCGACTTAACCTTAGGTTTTT). All miRNA inserts were cloned between HpaI and XhoI sites. Expression of GFP was under direction of the CMV promoter. For LMO4 overexpression experiments, the LMO4 open reading frame was cloned into pSicoR between NheI and AgeI sites, to be expressed as a GFP-LMO4 fusion protein. Lentivirus packaging was provided by System Biosciences (Palo Alto, CA). Titers of VSV-G pseudotyped viral particles were ~10^7^ IFUS/mL.

### Cortical cultures

Embryonic cortical cultures were prepared as previously described (103, 104). Briefly, e14.5 cortices were dissected and gently dissociated by papain digestion. A single cell suspension was prepared and plated onto poly-D-lysine (100μg/ml, Sigma) and laminin (20μg/ml, Life Technologies) coated coverslips in cortical culture medium. Cells were infected with lentivirus, and cultured on coverslips placed in 6-well plates for 7 days in growth media (50% DMEM, 50% neural basal media, supplemented with B27, BDNF, forskolin, insulin, transferrin, progesterone, putrescine, and sodium selenite). Under these conditions we observe >~95% neurons and very few (<~5%) astrocytes.

### Immunocytochemistry of cultured cells

Cells were fixed with 4% paraformaldehyde (PFA) in PBS. Coverslips were blocked with PBS containing 0.1% Triton-X100, 2% sheep serum, and 1% BSA, and cells were then incubated with primary antibodies against cell specific markers: anti-CTIP2 (AbCam, rat monoclonal, dilution 1:500), and/or anti-SATB2 (AbCam, rabbit monoclonal, dilution 1:100), and/or anti-LMO4 (AbCam, rabbit polyclonal, dilution 1:200). Following wash steps, the cells were incubated with secondary antibodies conjugated with fluorophores: anti-rat (Pierce, CY3-conjugate, dilution 1:1000), and/or anti-rabbit (Pierce, CY5-conjugate, dilution 1/1000), and anti-GFP (AbCam, goat-FITC conjugated, dilution 1:250). Cells were re-fixed in 4% PFA, and washed thoroughly in ddH2O. The coverslips were then mounted on microscope slides using Fluoroshield with DAPI (Sigma), and left overnight to dry. The following day, slides were sealed with clear nail polish, and imaged on a Zeiss AxioImager microscope. We counted neurons in 16 randomly selected high-powered fields, blinded to experimental condition. The experiments were replicated in n=5 independent cultures. Statistical analyses were carried out in R version 4.0.2 using a regression analysis taking all of the experiments (control/LOF/GOF/GOF+LMO4) into account. Specifically, we considered the scrambled control group as the reference level and defined dummy variables for miR 409-3p LOF/GOF/GOF+LMO4-rescue. We then regressed the outcome variable on the dummy variables, and then tested how each group differs from the scrambled control via Wald’s test. We also performed an ANOVA analysis under the regression framework to test whether there is an overall mean difference across the experiments. For the interaction analyses, we additionally defined a dummy variable for the LMO4 ORF overexpression experiment, and regressed the outcome variable on GOF, LMO4 and the interaction between GOF and LMO4. We tested whether the interaction coefficient is zero via Wald’s test.

### *In utero* electroporation

For miR-409-3p gain-of-function experiments, a CAG/H1 promoter plasmid was used to drive expression of tdTomato (tdT) and mature miR-409-3p. For control experiments, a similar plasmid was used to drive expression of tdT alone (control) or tdT and a scrambled control microRNA (scrambled control). *In utero* electroporation of embryonic mouse cortical neurons was carried out as previously described (15). Briefly, 750nL of plasmid DNA at 1μg/μL mixed with 0.005% Fast Green was injected into the lateral ventricle of e13.5 CD1 mouse embryos *in utero*. Plasmids were electroporated into the neocortical ventricular zone using 5mm diameter platinum disk electrodes and a square-wave electroporator (Nepa21, NepaGene) with five 28-Volt pulses of 50 milliseconds duration at 950 millisecond intervals, as previously described (105). Electroporated embryos were collected for analysis at e18.5.

### Immunocytochemistry of brain sections

Embryonic brains were drop-fixed in 4%PFA/PBS and post-fixed overnight in 4%PFA/PBS at 4°C, then equilibrated in 15% sucrose then 30% sucrose for cryopreservation over two days. Fixed and cryopreserved embryonic brains were sectioned on a sliding microtome at a thickness of 40 microns. Sections were wet-mounted onto microscope slides, allowed to dry for 2 hours at room temperature, fixed in 4% PFA for 10 minutes, then washed twice in PBS. Sections were incubated with primary antibodies overnight at 4°C, and appropriate secondary antibodies were selected. Antigen retrieval methods were required to expose the antigens for some of the primary antibodies; sections were incubated in 0.1M citric acid, pH 6.0 at 93-95°C for 10 minutes. Primary antibodies were used as follows: rat anti-Ctip2 (1:200, Abcam ab18465), rabbit anti-RFP (1:100, Rockland, 600-401-379). Immunocytochemistry was performed as previously described (26).

### Microscopy and image analysis

Entire embryonic hemispheres were imaged with fluorescence microscopy (Axioimager widefield fluorescence microscope, Zeiss) to evaluate location of electroporated cells and axonal projections. Individual cells were imaged with confocal microscopy (LSM710 Confocal, Zeiss). We counted neurons in 4 randomly selected confocal z-stacks, blinded to experimental condition. The percent of Ctip2+/tdT+ neurons was calculated. Experiments were replicated in n=3 biological replicates. Statistical analyses were carried out in Microsoft Excel using paired two-tailed t-tests with a significance threshold of p<0.05. We controlled for electroporation efficiency between the control and experimental groups by calculating the number of CTIP2+ cells as a percentage of tdT+ (transfected) cells.

## Data Availability

The differential miRNA expression data have been deposited in the GEO repository, accession number GSE116112.

---

**Supplemental Figure 1.**
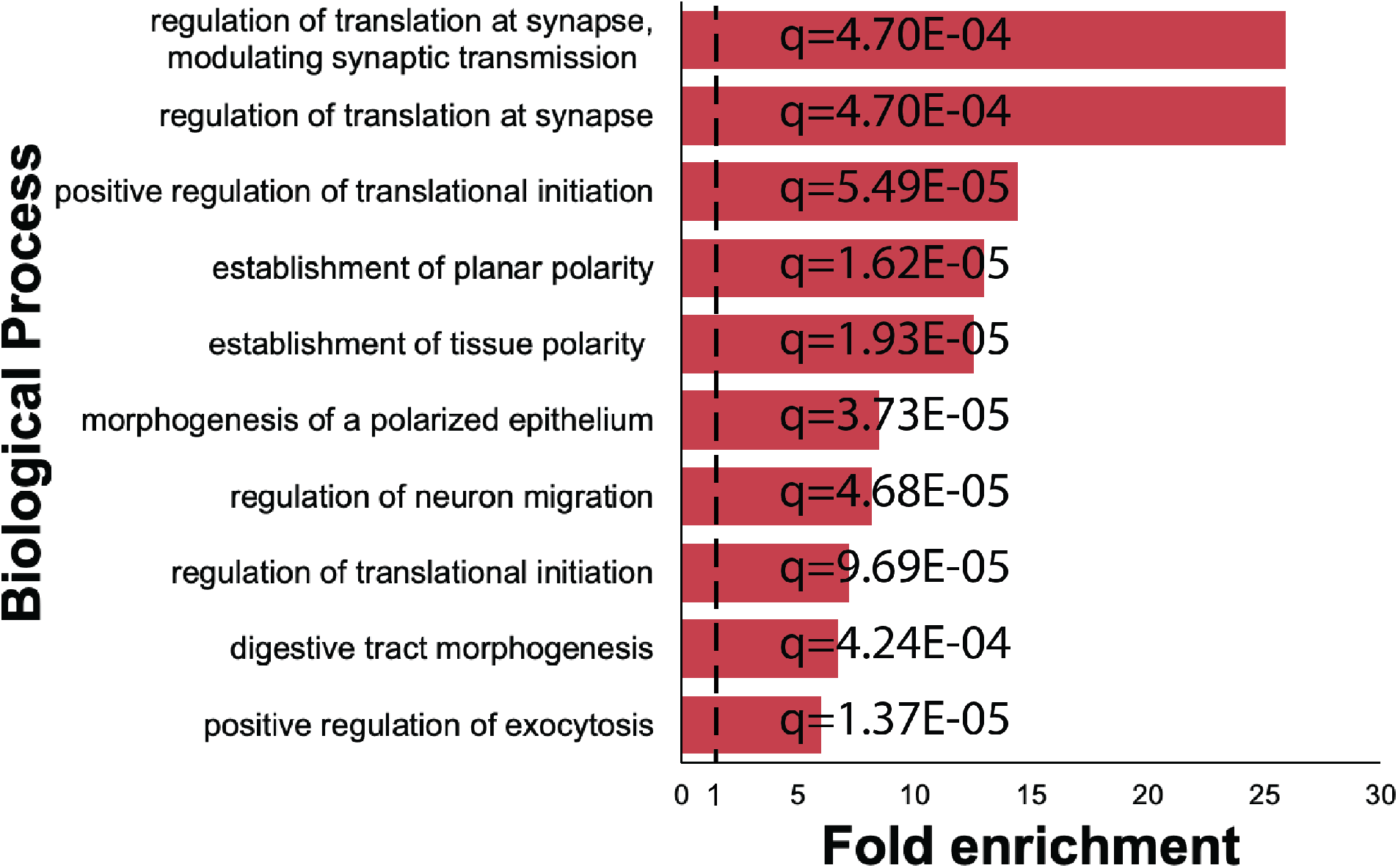
Gene Ontology (GO) analysis of predicted targets of miR-409-3p reveals over-representation of multiple biological processes relevant to cortical development. Biological process is along the y-axis, Fold enrichment is along the x-axis, and q value is given along the bars. The top ten over-represented processes are illustrated.

**Supplemental Figure 2.**
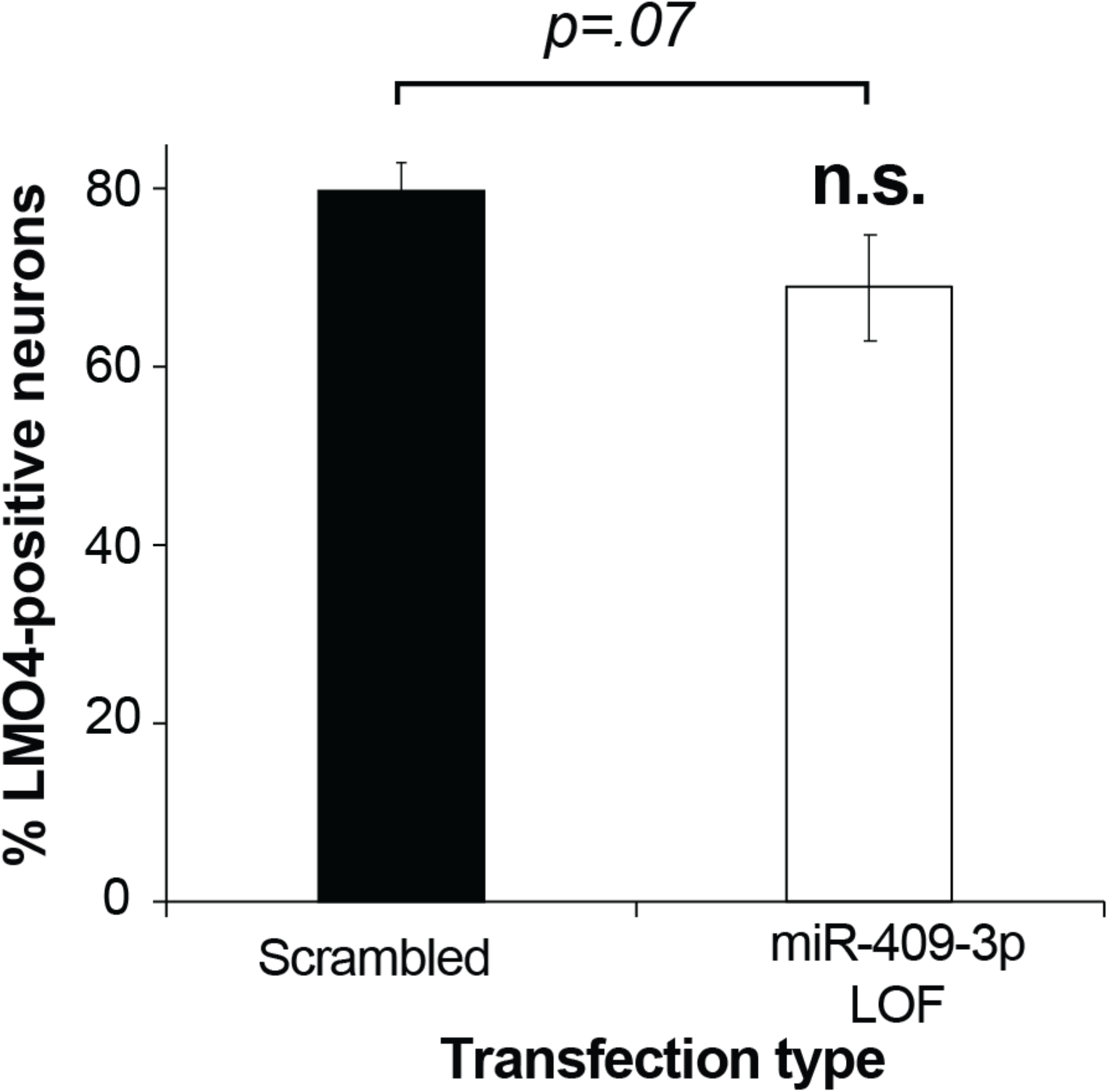
miR-409-3p loss of function does not alter endogenous LMO4 expression in cultured embryonic cortical neurons. miR-409-3p antisense loss-of-function (LOF) in cultured embryonic cortical neurons results in no significant change in expression of endogenous LMO4, compared to scrambled control, by immunocytochemical analysis. Error bars represent SEM. n.s. not statistically significant.

**Supplemental Figure 3.**
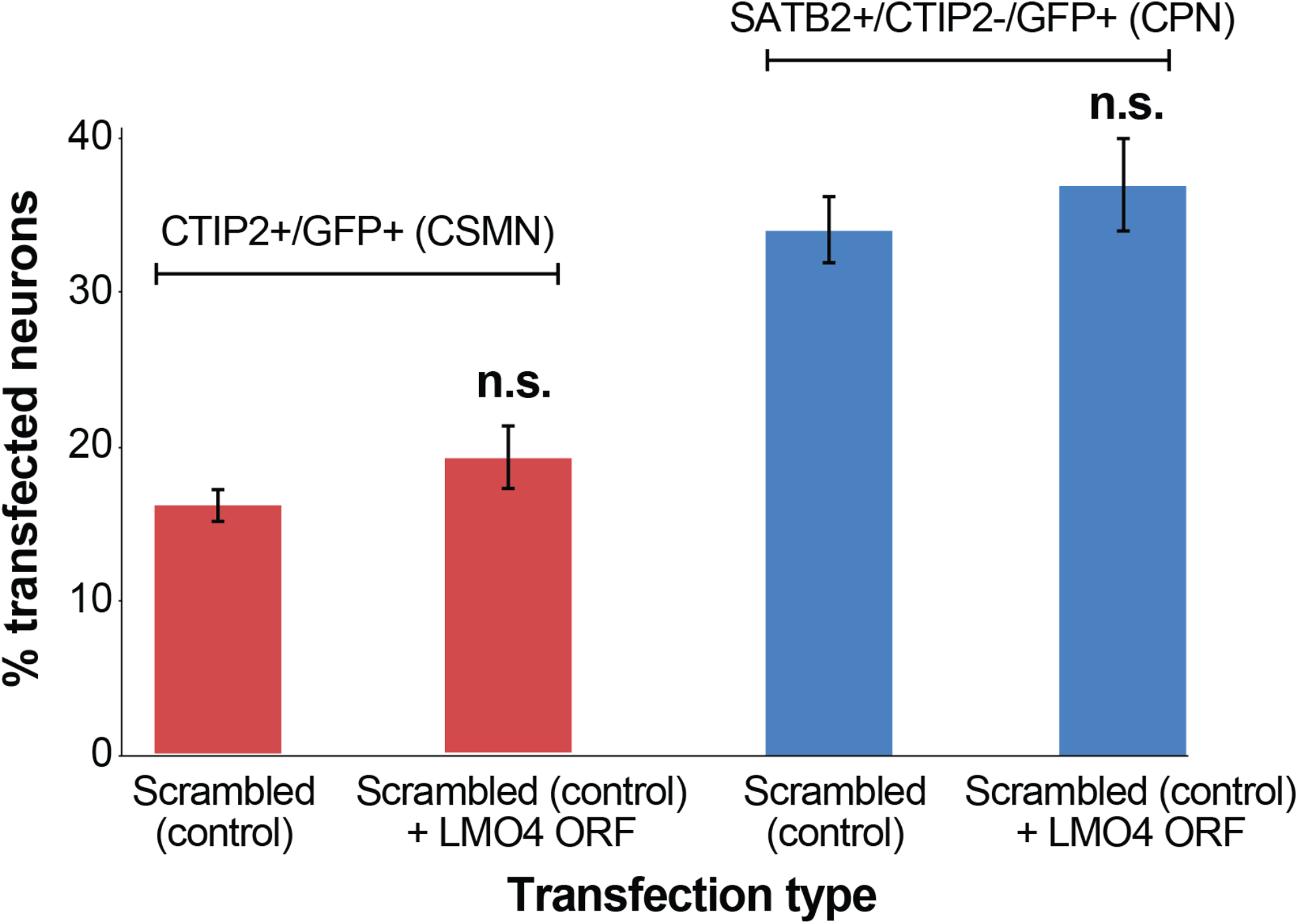
Overexpression of LMO4 ORF alone does not significantly affect CTIP2 or SATB2 expression in cultured embryonic cortical neurons. Overexpression of the LMO4 ORF in cultured embryonic cortical neurons results in no significant change in the percent CTIP2+/GFP+ neurons (CSMN), nor in the percent SATB2+/CTIP2−/GFP+ neurons (CPN), compared to control. Error bars represent SEM. *p<0.05; n.s. not statistically significant.

**Supplemental Table 1.**
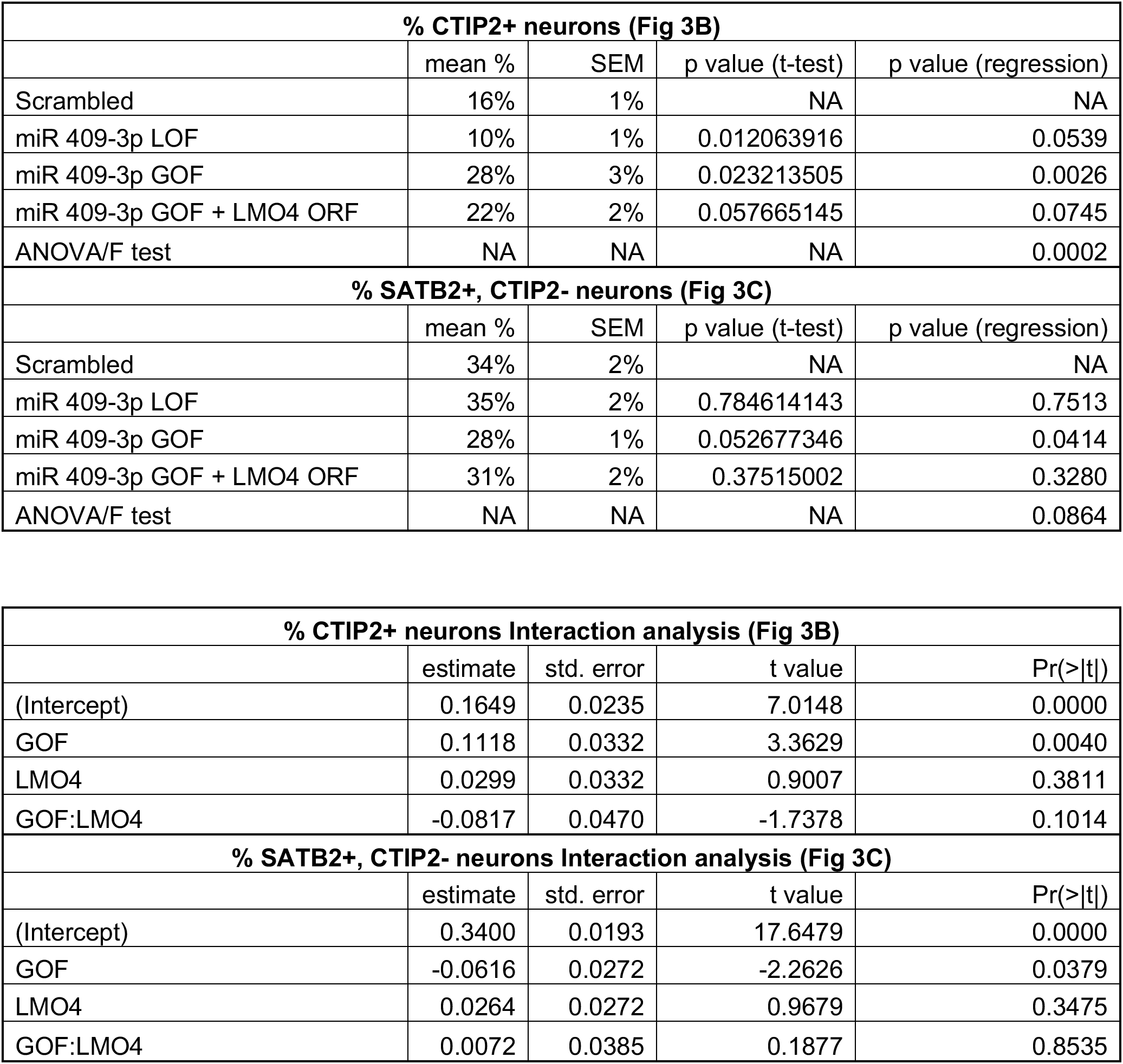
Summary of statistical analyses of miR-409-3p GOF and LOF studies *in vitro*. Results of individual t-tests, regression analysis, and ANOVA for miR-409-3p GOF and LOF. Regression automatically carries out an F test, which is equivalent to one-way ANOVA. Consistent with this, when we specifically ran an ANOVA, we obtained the same result as we did from the F test analysis in our regression analysis. Interaction analysis carried out to determine the effect of LMO4 ORF overexpression on miR-409-3p GOF.

